# Large complex structural rearrangements in human genomes harbor cryptic structures

**DOI:** 10.1101/2024.12.19.629504

**Authors:** Peter A. Audano, Carolyn Paisie, The Human Genome Structural Variation Consortium (HGSVC), Christine R. Beck

## Abstract

Structural variation is a major contributor to human diversity, adaptation, and disease. Simple structural variant (SV) types include deletions, insertions, duplications, inversions, and translocations, and SVs account for most of the variable bases between genomes. Complex structural variants (CSVs) that consist of one or more simple events *in cis* appear more frequently in diseases and cancers where DNA repair, apoptosis, and cell cycle checkpoints are compromised, although CSVs can also appear in germline genome sequences of healthy individuals. CSVs are often characterized by short tracts of homology or no homology, and while CSVs are more prevalent in complex regions that contain large repeats, smaller stretches of homology can also enable their formation across more unique loci. Long-read assemblies have increased the size of detectable SVs and expanded variant detection into more complex regions of the genome, and while they reconstruct CSVs, methods for identifying CSVs from assemblies is limited. Here, we have developed a new assembly-based approach to trace through complex loci rather than relying upon reference representations of alignments. We can now access CSVs in large complex segmental duplications, reveal structures that were previously unknown, and identify SV breakpoints with greater accuracy. We find 72 large CSVs per genome and 128 unique complex structures and CSVs in highly repetitive regions can now be detected including several distinct complex events in repetitive *NBPF* genes that was not previously callable with short-read or long-read CSV methods. This approach is implemented within a key assembly-based variant calling tool, PAV, and represents a substantial improvement identifying complex variants now ascertainable from contiguous genome assemblies.

## Introduction

A majority of SVs can be categorized as “simple”, comprising individual insertions, duplications, deletions, balanced inversions, and translocation events. CSVs, conversely, are broadly defined as combinations of simple SVs that arise in one single event and therefore comprise more than one repair junction. Like simple SVs, CSVs can also contribute to human diversity and disease including Mendelian disorders, autism, and cancer (Brand et al. 2015; Gu et al. 2015; Collins et al. 2017; Li et al. 2020; Wahlster et al. 2021).

While CSVs may be mediated by large segmental duplications (SDs) with highly identical repeat copies (1+ kbp, > 90% identity), they often lack features consistent with nonallelic homologous recombination (NAHR). Instead, replication-based double strand break repair mechanisms are often associated with CSVs. For example, shorter tracts of breakpoint homology are often observed at junctions of repair, which are consistent with microhomology-mediated break-induced replication (MMBIR) (Hastings et al. 2009). Blunt-ended junctions and other breakpoint features consistent with non-homologous end joining (NHEJ) (reviewed in Lieber (2010)) and microhomology-mediated end joining (MMEJ) (reviewed in McVey and Lee (2008)) are also often observed at CSV breakpoints, however complex events consisting of duplicated segments *in cis* with other simple events are not thought to occur solely by end joining mechanisms. For example, chromothripsis shatters chromosomes and joins fragments by NHEJ, however there are rarely duplicated segments in these rearrangements (Hu et al. 2024). Because some break repair pathways also utilize low-fidelity polymerases, single nucleotide and insertion-deletion variants are also sometimes observed *in cis* with CSVs (Beck et al. 2019).

Previous studies based largely on targeted analyses and short-reads have identified a number of CSV patterns in both normal individuals and affected individuals (Zhang et al. 2009; Brand et al. 2015; Collins et al. 2017; Collins et al. 2020; Li et al. 2020). While short-reads often rely on statistical clustering of copy-number events (Zhao et al. 2016; Zhou et al. 2024), long-read methods based on k-mers and machine learning have recently been developed (Lin et al. 2022).

Assemblies from long-reads now regularly span whole chromosomes, allowing the resolution of complex rearrangements without reference bias. We have observed that alignment tools, such as minimap2 (Li 2018; Li 2021), leave fragmented alignment patterns through CSVs leaving an opportunity for variant calling methods (**Fig. 1**). Redundancies in the alignments through repeats that mediate SVs map genome sequence to multiple locations obscuring the true structure of SVs (**Fig. 2A**). To properly detect large SVs, PAV removes redundancies (**Fig 2B**), which makes better breakpoint predictions, removes falsely duplicated bases, and eliminates artifacts such as false gene-fusion events apparent in the alignments (**Fig 2B**) (**Methods**). Other alignment artifacts around alignment breaks also produce complex substitutions, which PAV identifies with an alignment-free approach and removes (**Methods**). These steps are essential for increasing PAV’s sensitivity for large SVs and producing accurate callsets.

**Figure 1.**
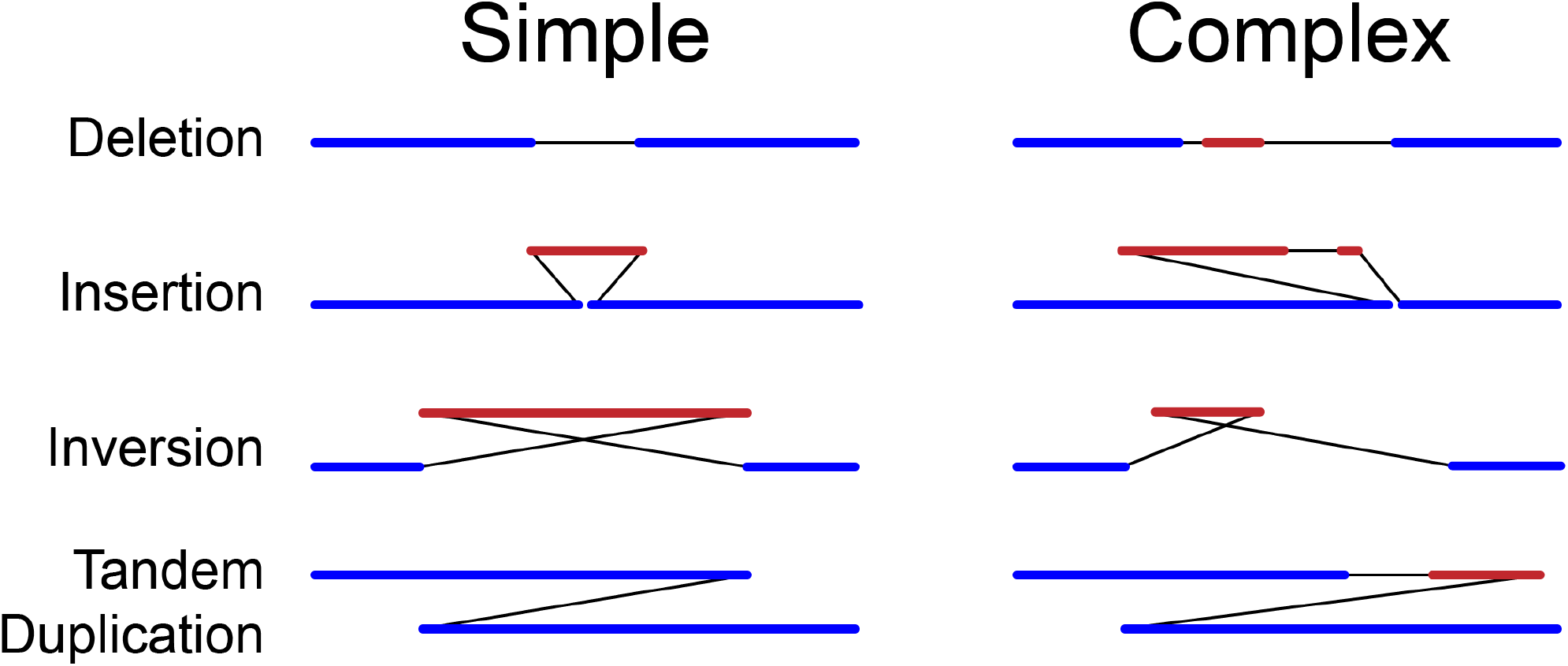
Choosing anchors for alignment-truncating SVs. Anchors (dark blue) after alignment trimming in query space (Trim-Qry) appear on each side of a large SV with segments of aligned or unaligned query sequence between anchors (red) and connections between them in assembly coordinates (black lines). PAV recognizes three types of simple SVs (left), deletions, insertions, inversions, and tandem duplications, each with a distinct alignment signature. For simple inversions, non-anchor query sequence (red) must be aligned in reverse orientation. All other variants fall into the complex category (right), which often have signatures resembling simple SVs. Not all simple variants will have exact breakpoints, for example an insertion with a small deletion-like gap at the breakpoint, and a scoring model is used to determine if these SVs fall into a simple SV category or the complex category.

**Figure 2.**
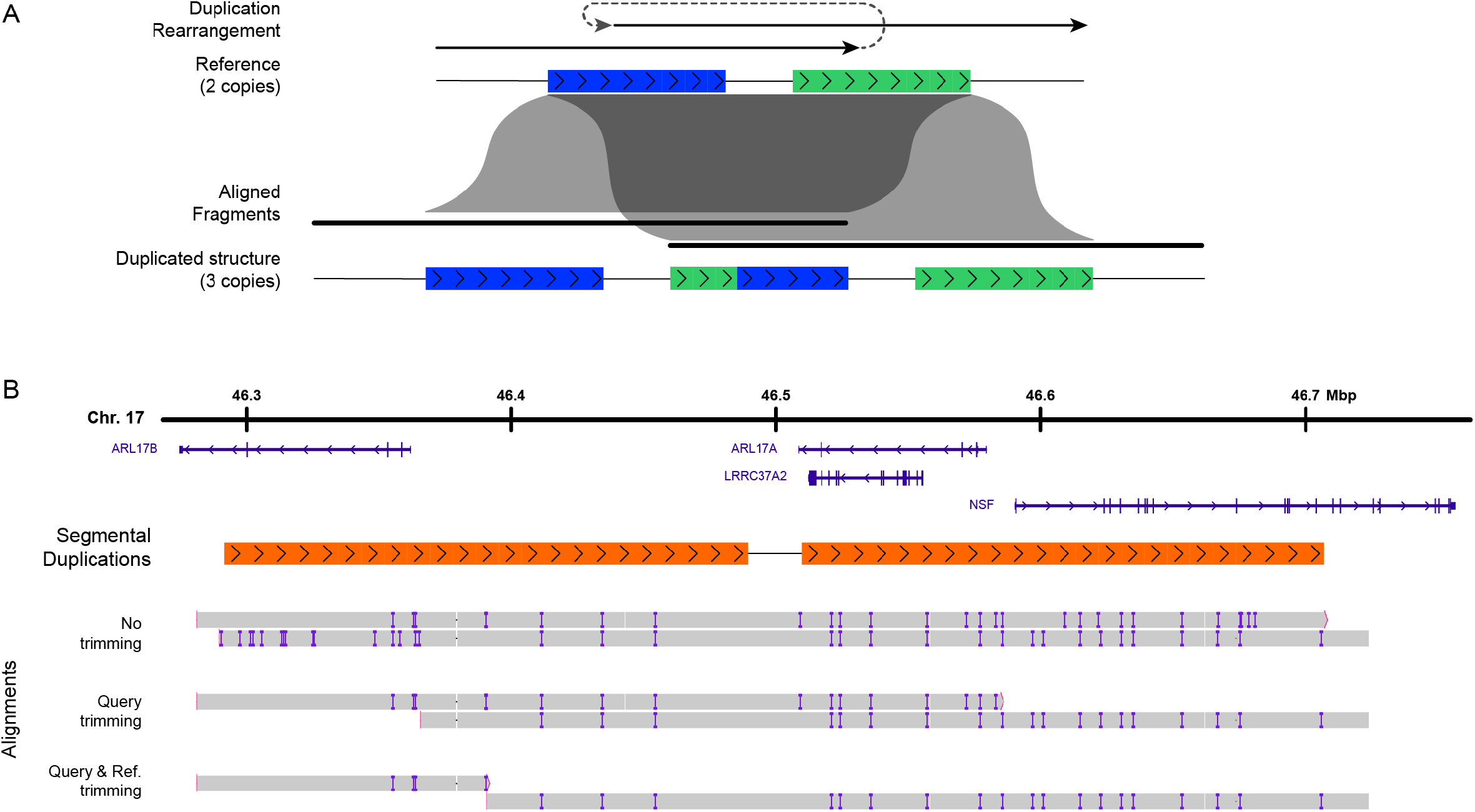
PAV alignment trimming example in an SD-mediated tandem duplication. **A)** A pair of SDs (blue and green) in the reference (Reference) recombines (Duplicated Rearrangement) resulting in a tandem duplication with three SD copies where the middle copy is a chimera of the original SDs (Duplicated Structure). The alignment fragments into two parts (Aligned Fragments) with the middle chimeric repeat aligned to both copies. **B)** An example alignment where a pair of reference SDs (orange, 197 kbp, >99% identical) mediated a tandem duplication represented in the assembly where the assembly has three SD copies. The initial alignments (“No trimming”) maps the middle SD copy to both reference copies it appear that four SD copies are present and suggesting an *ARRL17B*-*NSF* gene fusion. PAV first trims query bases from each end (“Query Trimming”) maximizing the number of SNVs (not shown), insertions (purple bars), and deletions (gaps in gray bars) that are removed, which reveals a more accurate representation of the duplication and a more precise location for duplication breakpoints in both reference and query coordinates. After trimming, the false *ARRL17B*-*NSF* gene-fusion is no present. PAV’s large variant discovery process primarily uses this alignment representation. To choose a reference location for insertions, a second stage further trims against redundantly-aligned reference bases (“Query & Ref. trimming”). The UCSC genome browser was used as a guide, and alignment representation was generated by the Integrative Genome Browser (IGV). Alignments and annotations are in GRCh38 coordinates. Psedogenes and RNA genes are not shown.

Building on PAV’s proven ability to handle large SVs (Ebert et al. 2021; Logsdon et al. 2024), we developed a new approach that uses a unique assembly-centric method of tracing alignments through both reference and query (assembly) sequences to reconstruct both simple SVs and CSVs. We have now incorporated this method into a beta version of our variant caller, PAV, to allow comprehensive variant detection in one tool.

## Results

### Complex callset production

As part of the Human Genome Structural Variation Consortium (HGSVC), we developed 65 phased human genome assemblies (130 phased haplotypes) and a matching variant callset identifying single nucleotide variant (SNVs), insertion/deletion variants (indels, < 50 bp), and SVs (≥ 50bp) using PAV with a large number of callers for orthogonal support (Logsdon et al. 2024). In this paper, we expand the callset from these assemblies using a new version of PAV (3.0) to identify CSVs.

Utilizing trimmed alignments, PAV calls CSVs by tracing breakpoints through both reference and the assembly sequence (**Methods**), which we applied across two references, GRCh38-NoALT (GRCh38) and T2T-CHM13v2.0 (T2T-CHM13) (**Methods**). For this effort, we exclude centromere and satellite repeats where alignments are less reliable (Ebert et al. 2021; Logsdon et al. 2024), eliminate assembly artifacts, and create a nonredundant CSV callset (**Methods**).

### PAV identifies large CSVs

When our assemblies are aligned to the T2T-CHM13 reference, we identify an average of 72 CSVs per sample (51 – 91 per sample, or 26 – 51 per assembled haplotype) (**Tables S1**,**S2**). Combining all events across the 130 phased haplotypes, we find a total of 5,122 CSVs, which merge to 1,247 nonredundant CSVs (**Table S3**). We replicated our analysis with GRCh38 as the reference, and find a total of 9,255 CSVs, which merge to 1,524 nonredundant CSVs (**Table S3**). Most CSVs are large with median CSV sizes of 91.5 kbp (T2T-CHM13) and 101 kbp (GRCh38). As has been previously shown with simple SVs, we identify more CSVs per sample with GRCh38 as the reference (120 GRCh38 vs 72.0 T2T-CHM13 on average), suggesting that GRCh38 contains complex structural errors in addition to simple structural errors (Aganezov et al. 2022; Logsdon et al. 2024).

We identify 128 distinct CSV structures in T2T-CHM13 (**Table S4**) and 187 in GRCh38 (**Table S5**). Common structures in T2T-CHM13 include INVDUP-INV-DEL (174 CSVs), DEL-INV-DEL (34 CSVs), DUP-NML-DUP (12 CSVs), and INVDUP-INV-INVDUP (8 CSVs) where DEL is deleted sequence, INV is inverted sequence (but not duplicated), INVDUP is sequence that was duplicated and inverted, and NML is a segment that was not deleted, inverted, or duplicated within the CSV (**Fig. 3**). While most CSVs have simple duplications or inverted duplications, others create higher-copy segments including DUP-TRP-DUP (6 CSVs) where TRP is a triplication (**Fig. 3**).

**Figure 3.**
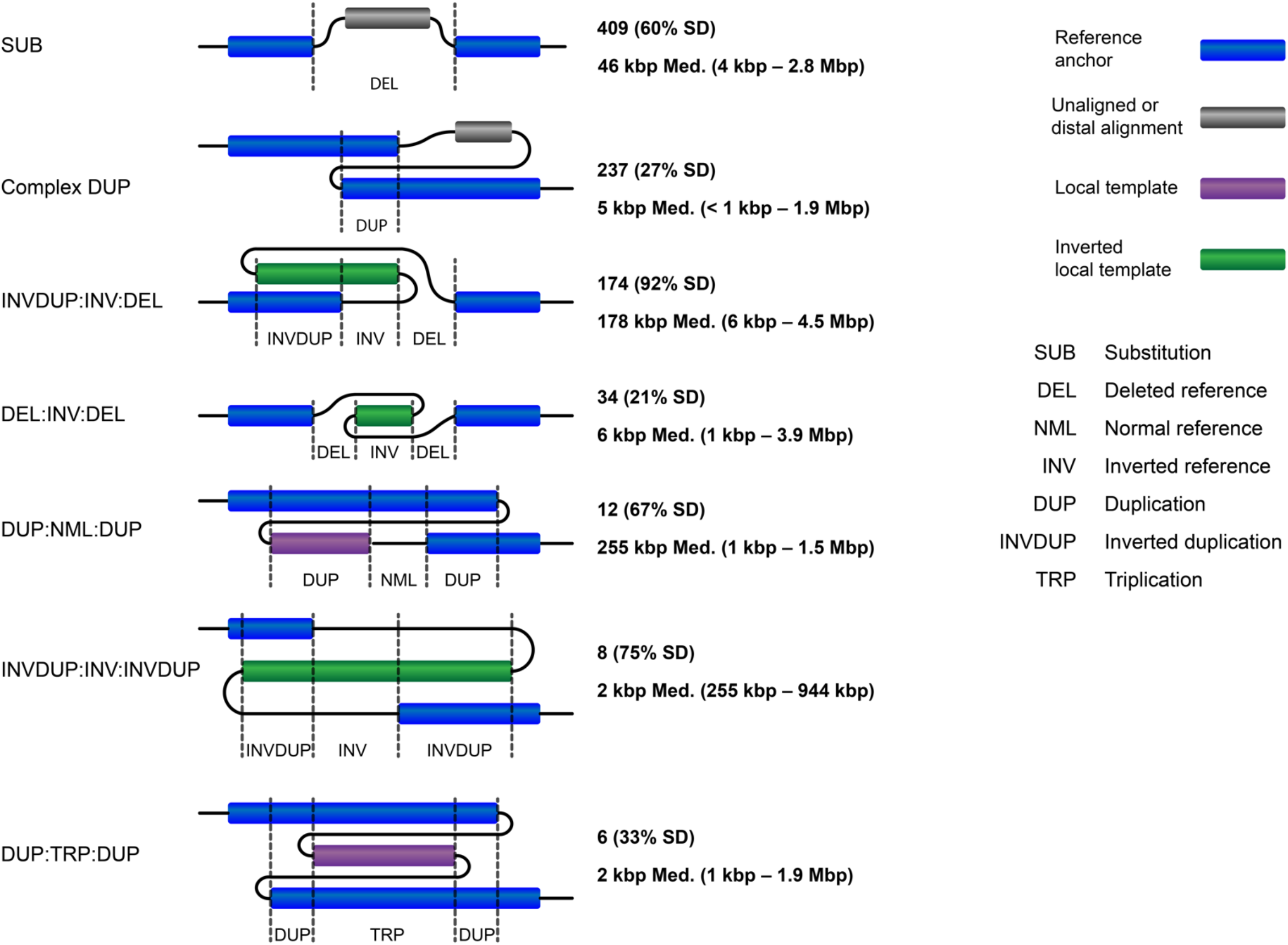
Template switches across CSV types generate different local structures at the SV site. Each SV is anchored by alignments on each end (blue bars). Many CSVs have unaligned sequences embedded within them (gray bars), which do not contribute to the local structure. When templates are derived from sites proximal to the SV site, it creates a distinct local structure composed templates replicated in reference orientation (violet bars) and inverted orientation (green bars).

**Figure 4.**
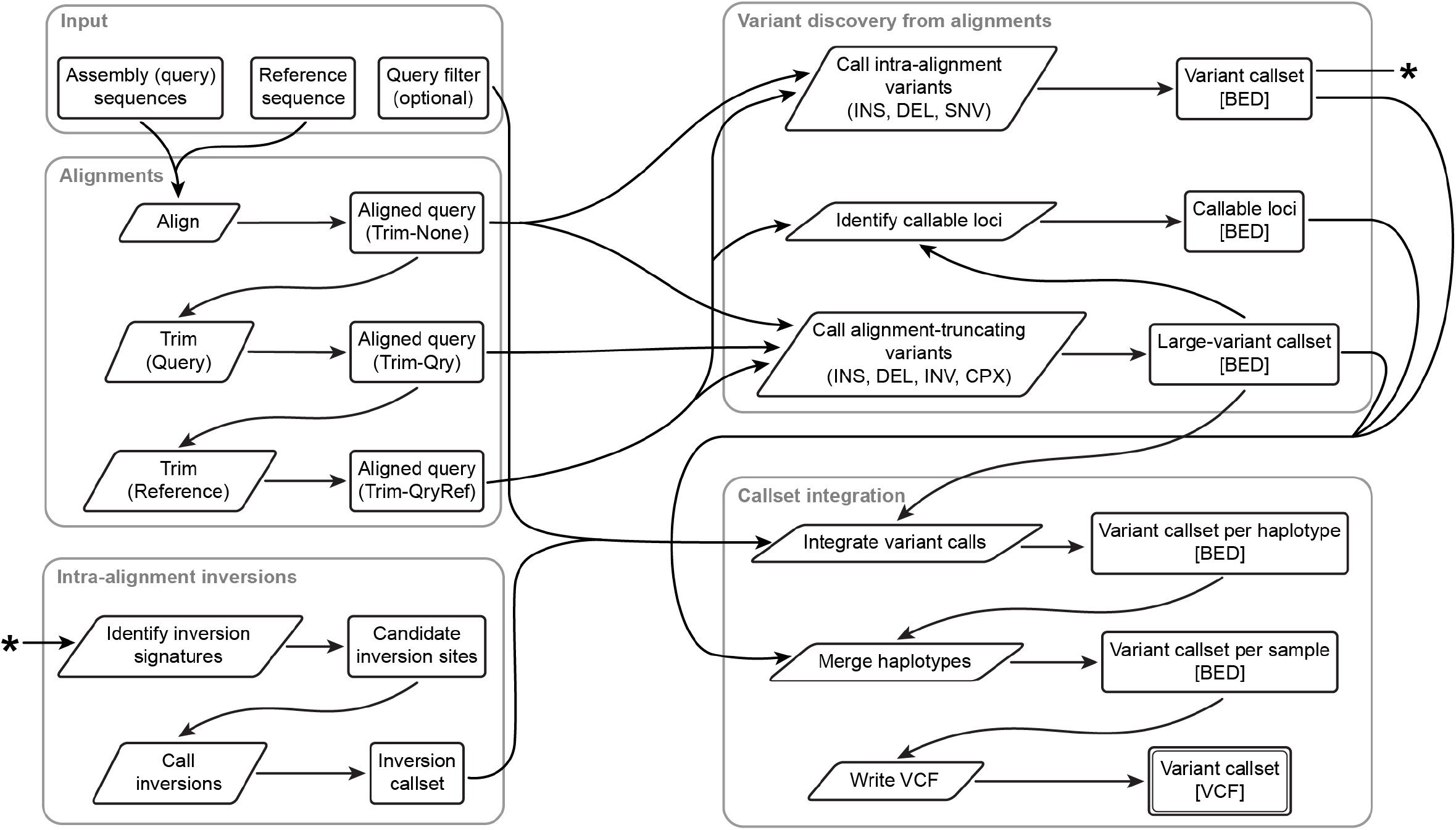
PAV pipeline flowchart. The PAV variant calling process is implemented in several distinct steps taking input through alignments, variant discovery, and callset integration. In the alignment steps, assemblies are aligned to a reference genome and several stages of alignment trimming are applied. Variant discovery steps use evidence from alignments to make variant calls including intra-alignment calls using alignment CIGAR strings and variants that cause alignment records to fragment into pieces (“alignment-truncating” variants). A separate step calls intra-alignment inversions where alignments were pushed through inverted loci embedding signatures of inversions in CIGAR strings (matched INS/DEL SVs and clusters of SNVs and indels), and it attempts to identify inversions at those sites. The asterisk indicates that inversion signatures use the intra-alignment variant callest. Callset integration takes PAV output from all stages and all haplotypes and integrates it into a final callset. Arrows are not shown for all stages that use the input query sequence, for example, most variant calling steps extract sequence from the query.

### Many CSVs have a simple local structure

When CSV breakpoints are clustered in one location, a complex structure is apparent, for example, INVDUP-INV-DEL. However, other CSVs have distal duplications or unaligned sequence obscuring the complex structure. These often appear as substitutions or tandem duplications with additional sequence between the breakpoints (**Fig. 3**). In our callset, we identify 409 substitution CSVs and 237 duplication CSVs, and while alignments suggest deletions or tandem duplications, PAV identifies the complex unaligned or distally-aligned sequence embedded within them by tracing the reference and assembly through breakpoints (**Fig. 3**) (**Table S4**).

Of these complex substitutions, 92% were linked with an unaligned sequence (median 13.0 kbp), and 23% were found with up to 6 templates distal to a reference deletion (median 59.7 kbp). Similarly, 97% of complex tandem duplications were linked with unaligned sequence (median 1.0 kbp), and 32% were found with up to 9 templated sites distal to the tandem duplication (median 38.4 kbp). While these substitution and duplication CSVs do not create unique complex structures at the SV site, PAV can now link 22.6 Mbp of distally duplicated sequence and 17.2 Mbp of unmapped sequence to sites that appear to be simple deletion or tandem duplications. More precise alignment methods will be needed to map these unaligned complex sequences in reference genomes.

### Complex rearrangements in *NBPF* genes

At the *NBPF8* gene locus (chromosome arm 1p) in T2T-CHM13, we find three haplotype structures. A reference allele is observed with 13.7% allele frequency (AF). We find a DEL-INV-DEL CSV inverting *NBPF8* with 35.9% AF (Logsdon et al. 2024). Although this CSV merged into two distinct variant calls, it is unclear if this is due to merging error from polymorphisms within the CSV sequence or if it is evidence of recursion. A second CSV deletes *NBPF8* and replaces it with a duplicated copy of *NBPF9* (chromosome arm 1q) templated from 25 Mbp upstream (Logsdon et al. 2024). This event is present with 50.8% AF and averages 513 kbp. When haplotypes with the same CSV are compared, we find the CSV sequence between haplotypes averages from 91.6% to 96.3%. When haplotypes carrying different CSVs are compared, the CSV sequence identity averages from 70.7% to 89.6% suggesting that these are distinct CSVs and not alignment artifacts between similar SDs (**Table S6**). To confirm merge assignments, we applied spectral clustering across pairwise variant identities (**Methods**) and find that the *NBPF9* alleles associate with one cluster (**Table S7**). While CSVs are less likely to be recurrent (Beck et al. 2019), large repeats also mediate similar but distinct CSVs at the same site.

## Discussion

Genomes are subject to many mutational mechanisms that lead to base changes, small indels, and structural variation, all of which are important substrates for selection (Genomes Project et al. 2012; Sudmant et al. 2015; Collins et al. 2020; Ebert et al. 2021) as well as drivers of diseases (Beck et al. 2019; Li et al. 2020; Wahlster et al. 2021). In recent years, thousands of CSVs have been identified using abundant short-read Illumina genomes and statistical methods, which has been applied to both healthy and disease cohorts (Zhang et al. 2009; Brand et al. 2015; Collins et al. 2017; Collins et al. 2020; Li et al. 2020). Although long-reads can capture whole CSV structures in single read or assembly sequences, methods to resolve them have been limited.

By utilizing the full sequence of CSVs, we introduced a method to trace rearrangements through multiple breakpoints to reconstruct them to basepair accuracy. We leverage unique abilities of PAV to resolve redundant alignments through the repeats that mediate SVs including mobile elements and SDs and to identify and remove alignment artifacts that would otherwise lead to false CSV calls. We find a surprising number of CSVs appear to be simple substitutions or tandem duplications, although with large sequences either duplicated from a distal site or left unaligned. While PAV can now trace through alignments with greater accuracy to resolve CSVs, alignment improvements are needed to completely resolve many CSV events and refine their breakpoints.

The *NBPF* gene family has been subject to recent expansions and structural rearrangements with many copies present throughout the genome (Vandepoele et al. 2005; Jiang et al. 2007). The combination of long-read technologies and modern assemblies can now resolve the *NBPF8* locus routinely in all our haplotype assemblies (Logsdon et al. 2024) representing a significant improvement in recent years (Ebert et al. 2021). In addition to assemblies, this locus has evaded CSV detection for both short-read (Collins et al. 2020) and long-read methods (Lin et al. 2022). With assemblies now approaching T2T standards (Nurk et al. 2022; Logsdon et al. 2024), PAV’s CSV algorithm can now access large CSVs in rearrangement hotspots.

We share a CSV callset utilizing 65 samples and 130 phased haplotypes derived from those samples. As more long-read genomes are produced at increasing frequency for both healthy (Liao et al. 2023; Logsdon et al. 2024; Mahmoud et al. 2024) and patient (Cohen et al. 2022; Hiatt et al. 2024) cohorts, additional complex variation may be discovered and associated with disease or other phenotypic effects. By completely sequence-resolving these sites and identifying the true breakpoints, better predictions about the mechanisms of CSV formation and the effects of CSVs can be made, for example, identifying gene fusions (Wahlster et al. 2021) or determining if duplicated genes are likely functional. These advances in sequence data and methods now make it possible to study human genome variation with greater accuracy and identify both the effects of CSVs as well as the mechanisms that mediate them. To meet the demand of long-reads in modern genomics and the growing number of projects that depend on them, efficient and accurate tools such as PAV are needed to produce accurate callsets from small SNVs and indels to large SVs and CSVs.

## Software availability

A beta version of PAV described by this paper is now available (https://github.com/BeckLaboratory/pav).

## Methods

### Sample assemblies, genome references, and alignments

#### Samples

We utilized 65 phased assemblies (130 haplotypes) derived from 1000 Genomes samples that were generated by the Human Genome Structural Variation Consortium (HGSVC) (Logsdon et al. 2024). Briefly, the cell lines were sequenced with PacBio HiFi, Oxford Nanopore Technologies (ONT), and Strand-seq, and resulting data were used to produce near-T2T quality phased assemblies.

#### References

Variant discovery utilized two reference sequences, T2T-CHM13 (Nurk et al. 2022) and GRCh38 (Schneider et al. 2017) with ALT and patch sequences removed from GRCh38 (Ebert et al. 2021).

#### Alignments

Assemblies are aligned to each reference with minimap2 (2.28) (Li 2018; Li 2021) with parameters “-x asm20”. One independent callset is produced for each reference.

### A model for scoring alignments and variants

*Score model*. A multi-affine score model is used with default parameters matching the default minimap2 (Li 2018) double-affine alignment parameters. This model scores matches, mismatches, and gaps (insertions and deletions) using gap-open and gap-extend penalties for alignments and variant calls. PAV’s implementation accepts any number of gap-open/gap-extend pairs (multi-affine), and when scoring gaps, will try all and choose the lowest penalty. To support complex variant calls, the model also includes a template switch penalty, which by default is set to the cost of two 50 bp gaps under the affine parameters.

To apply these scores to alignments and variants, we define score constants R and a score function *score(o,l)* taking an operation code (o) and an operation length (l) and returning the score for that operation. Operation codes are defined to match CIGAR string operators (ignoring clipping and padding operations) with an additional element for template switches (o = ***t***), which has no representation in real CIGAR strings. For convenience, we define *gap(l)* as the cost of a gap (insertion or deletion) of length *l*.

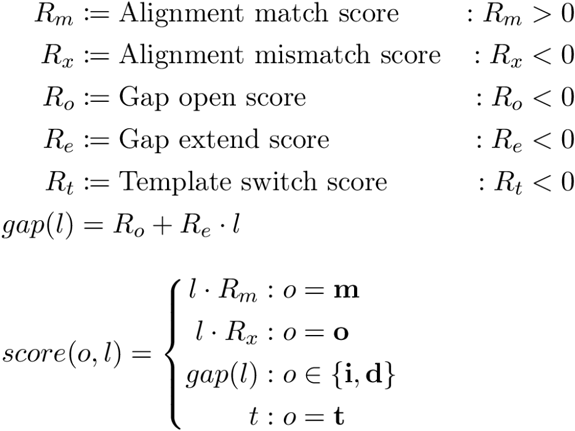

Regardless of the parameters used by the alignment tool to map queries to references, PAV uses one score model across the whole callset process. Each alignment record is scored by this model, and the same model is used again during large SV detection.

#### Scoring alignments

After each query alignment or trimming step, each alignment record scored by its CIGAR string (S(a_i_)). For convenience, we define cigar(a_i_) as a function that returns tuples of CIGAR operation codes and lengths for alignment a_i_.

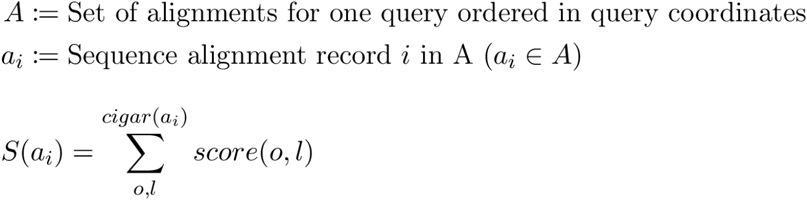

### Alignment trimming and aggregation

#### Motivation

When an alignment reaches a large difference between the query and reference sequences, it often stops aligning at that site and resumes alignments at another site, which fragments the alignment into multiple records. This is often observed around highly complex and divergent loci as well as simple SVs, for example, a large deletion might cause the alignment to terminate at one end and resume at the other. Proper handling of these alignment-truncating events is essential for identifying large variants. The threshold at which the alignment program decides to break the alignment is dependent on the tool and the alignment parameters. When SVs do not cause alignments to break, the SV is embedded in the CIGAR string of a single alignment record. To increase sensitivity and support a variety of alignment parameters, PAV includes methods for calling variants both within alignments and at alignment-truncated sites.

When alignments truncate and restart, query bases are often aligned on both ends. For example, the alignments around a 217 kbp tandem duplication (TD) mediated by SDs has three SD copies, but the initial alignments appear to have four copies and a gene fusion event (**Fig. 2**). After redundantly-aligned query bases are resolved, three SD copies are evident, the breakpoint locations of the TD event are more precise, and no gene fusion is suggested by the alignments (**Fig. 2**).

#### Trim stages

The initial alignments produced by PAV (Trim-None) have no alignment trimming applied. PAV first trims in query space only (Trim-Qry) such that no query base appears in more than one alignment record (i.e. each query base is aligned zero or one times). PAV performs a second round of trimming to remove redundancies in reference space (Trim-QryRef). These alignments are used for determining the breakpoint for insertion sequences mediated by repeats and for choosing the best paralog for variant calls in regions where more than one query aligns.

These trimming steps are used by downstream steps (**Fig. 5**).

**Figure 5.**
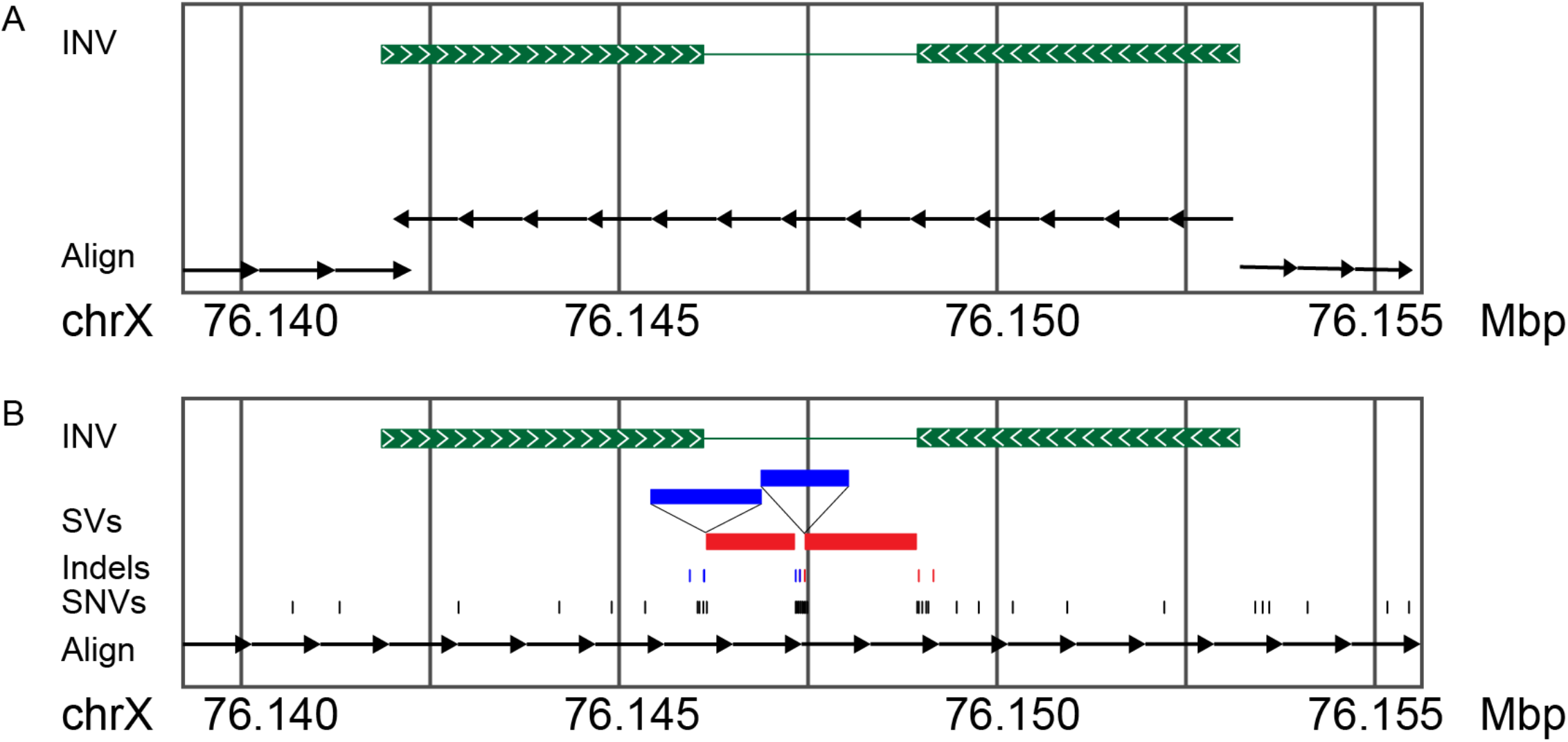
Alignments supporting inversions may be obscure. An example of two types of alignment signatures through inverted loci. In this example, inversions are mediated by inverted repeats (green). Alignment records and their relative orientation with the reference is shown (arrows) (A) Larger inversions cause alignment records through the inverted site to fragment and create an inversion signature composed of flanking records in the same orientation and the inverted record in the opposite orientation. These inversions are easily identifiable from the alignment pattern. (B) Smaller inversions may not cause the alignment to truncate, instead, the alignment tool pushes the alignment through the inversion leaving false SVs (long red and blue bars), clusters of false indels (small red and blue bars), and clusters of false SNVs (black bars). PAV recognizes these signatures and resolves the true inversion at these sites.

#### Algorithm

Two alignment records with overlaps in query coordinates (i.e. same query bases aligned in both records) are input into the algorithm. PAV generates a list of alignment events through all redundantly aligned bases on both records. It sets an initial cut site at the head of one list and the tail of the other (i.e. removing all redundantly aligned bases by maximally cutting one side and not cutting the other side). Cut sites are iteratively shifted through alignment events in both lists (going up one list and down the other) under two constraints, (a) chosen cut sites must not allow any redundantly aligned bases, and (b) if leaving unaligned query bases is unavoidable, it is minimized. The algorithm proceeds iteratively shifting the cut site in one list by one event, shifting the cut site on the other list to match these constraints, and scoring the number of indels and SNVs removed at each pair of cut sites. The cut-site location that maximizes the number of indel and SNVs discarded is chosen. If there is ambiguity, cut sites are chosen to preserve left-alignment conventions (Audano and Beck 2024). These constraints and the iterative algorithm reduce the search from O(n^2^) over all possible pairs of cut sites between the two lists to O(n). The same algorithm is applied to reduce Trim-Qry to Trim-QryRef by targeting and removing redundantly aligned reference bases.

#### Use and filtering

PAV utilizes all three trimming stages. Initial intra-alignment variants called from CIGAR strings are performed Trim-None alignments, and variants that are removed in Trim-QryRef alignments are marked with a “TRIM” filter in PAV’s output. Large alignment-truncating variant detection is done primarily with Trim-Qry alignments, however, Trim-QryRef is used to select an insertion site for SV duplications.

#### Alignment aggregation

Alignments sometimes fragment over arbitrary sites leaving deleted bases with paired insertions that match the deletion (i.e. no real variant) or break over small variants. For optimal PAV performance, small variants should be embedded within single alignment records (i.e. the CIGAR string) leaving large variants for the algorithm over alignment-truncating SVs. If not aggregated, these alignment breaks obscure true large variant signatures, for example, breaking a large duplication into many smaller structures and making complex SVs appear to be more complex than they are.

To prepare alignments for large variant calling, alignment records adjacent in query space are that are approximately colinear are aggregated. Colinearity is considered if the reference gap is 0 bp or greater (i.e. do not align to the same reference bases), otherwise, aggregation is not attempted. A score is computed by summing a gap penalty for the length of skipped reference bases (DEL), the length of unaligned query bases (INS), and a colinearity penalty computed as the absolute difference between the lengths of INS and DEL (i.e. 4,000 bp INS and a 3,000 bp DEL incurs an additional penalty equal to a 1,000 bp gap). If the summation of these gaps is less than a set threshold (the score of a 10 kbp insertion or deletion by default), then the two alignment records are joined into one at the junction. PAV’s alignment QC code is used to check the resulting alignment record for inconsistencies, for example, ensuring end locations agree with start positions and CIGAR operations in reference and query space after the alignment record is aggregated into one. The new alignment record is also re-scored using the scoring model. Aggregation is applied after query trimming and was performed for al Trim-Qry and Trim-QryRef alignments.

### Intra-alignment inversions

#### Introduction

Inversions identifiable to PAV generate one of two alignment patterns. A majority of detectable inversions aligned through minimap2 (Li 2018; Li 2021) cause the alignment to split over the inversion site (**Fig. 5A**), and these are easily detected by PAV and other assembly-based variant callers including DipCall (Li et al. 2018) and SVIM-asm (Heller and Vingron 2021). When alignments are not fragmented, they leave alignment patterns of false SVs, indels, and SNVs (**Fig. 5B**), which require a different approach for calling. In the final VCF, PAV reports the resolved inversion and the false SVs, indels, and SNVs are filtered (FILTER = COMPOUND) and linked to the inversion call (INFO/COMPOUND is the inversion ID).

#### Intra-alignment inversion signatures

Candidate inversion sites are identified by searching for inversion signatures. Clusters of indels or SNVs are found by merging SNVs and indels within 200 bp. Clusters must reach 500 bp in length (from the first indel or SNV to the last) and reach 10 events for indel clusters or 20 events for SNV clusters. Matched events (a deletion and an insertion in close proximity) are searched in a region 2× the SV length surrounding the event for a matching variant.

#### Convolution method

PAV determines the presence of an inversion with an alignment-free method k-mer densities. A reference site around the inversion signature is chosen and the corresponding query sequence aligned to the region is found by lifting reference coordinates through the query alignments to the query sequence. Both the reference and matching query sequence is extracted, both in reference orientation. The reference sequence is translated to a set of k-mers (31-mer by default). High-count k-mers are identified by summing the number of occurrences of a k-mer and it’s reverse-complement and removing both if the sum exceeds a threshold (10 by default). For k-mers that are not removed, they are retained in the inversion set in their original orientations. The query sequence is translated to a list of k-mers of the same size (31-mer by default) and all k-mers that do not match a reference k-mer in either orientation is removed from the list, which prevents SVs inside inversions from obscuring the inversion structure. The list of k-mers is then translated into three lists of true/false values (1 or 0, respectively): one list for reference matches in forward-only orientation, one for forward and reverse orientation, and one for reverse-only orientation. These three lists are predictors for inverted loci (reverse-only sites) and their inverted repeats (forward and reverse sites).

Smoothing is performed over each of these three states. Previous versions of PAV used a kernel density estimation (KDE), which was rapidly degraded in performance over larger inversions because the algorithm is O(n^2^) in the length of inversion sites (compute the sum of n kernels each of n k-mer sites). By sparsely estimating density and filling in sites where density changed rapidly, PAV was able to reduce some computational burden, although performance was still significantly degraded.

PAV now replaces the KDE method with a convolutional method. First, a kernel is chosen using a truncated normal distribution with bandwidth 100.0 truncated at Z = ± 3.0 such that the sum of probabilities from Z = -3.0 to Z = 3.0 being 1.0, as defined for truncated normal distributions. These parameters cover 300 sites upstream and 300 bp downstream of a site, including the site itself (601 position window) where weights decreasing with distance from the center and the sum of weights is 1.0, consistent with truncated normal distributions. This distribution is then centered at each position in a boolean list, and the weighted sum is computed for the center-most site. When a site is surrounded by the same state for 300 bp in either direction, the sum is approximately 1.0 (with small error allowed for floating-point precision), and it decreases as the purity of the state around the site decreases where distal sites are weighted less heavily than proximal ones within the window. To preserve a maximum of 1.0, the sequence window is expanded by half the convolutional window, which is removed again after computing the convolution, otherwise, the flanking sites would reach a maximum of 0.5. Site expansion is not strictly needed for PAV’s algorithm, but it makes it easier to interpret results directly from the density tables if needed. Since this approach is bound by O(n^2^) if the window is large, we utilize a Fast Fourier Transform (FFT) by default to reduce runtime to O(n × log(n)).

The convolution for each list of states is performed (forward-only, forward and reverse, and reverse-only). We then assign a smoothed state to each site by taking the maximum of the three states for each site. Two patterns are then searched for inversions, one with inverted repeats on each side and one without. Inversion breakpoints are set at the boundaries of the inner-most reverse-only site, and if found, outer-breakpoints are reported with the inversion at the outer-most forward-and-reverse sites. Some inverted sites contain repeats and do not have single transitions through these states (e.g. forward -> forward+reverse -> reverse -> forward+reverse -> forward), and for these, state transitions are chosen to balance forward+reverse sites on both sides (i.e. most similar sizes).

### Alignment truncating variants

#### Introduction

Large SVs break alignments into multiple records at SV sites (**Fig. 1**). These SVs can have simple or complex structures, and they can be identified by tracing the query sequence through alignment records. To achieve correct breakpoints for repeat-mediated SVs, query-trimmed alignments (Trim-Qry) are used for this step (**Fig. 2**). PAV requires all alignment-truncating variants to be anchored on each end with an SV sequence between anchors (**Fig. 1**).

When considering complex SVs, choosing anchors is not trivial. For example, a complex deletion with a small aligned fragment in the middle could be two simple deletions or a complex deletion with a small templated insertion in the middle (**Fig. 1, complex deletion**). PAV will need to consider all possible representations of variants through alignments and choose an optimal one.

#### Anchor candidates

Large variants are anchored on each side by alignment records often spanning into sequence uniquely mappable to a single locus in the reference genome. A pair of anchors for a large SV must satisfy two criteria, (a) They must be colinear (same query sequence aligned in the same orientation), and (b) they must have a sufficiently high alignment score to support a variant call between them. PAV identifies pairs of anchor candidates using an exhaustive search over all possible anchor pairs.

The following algorithm describes building a set of candidate anchor pairs (*I*).

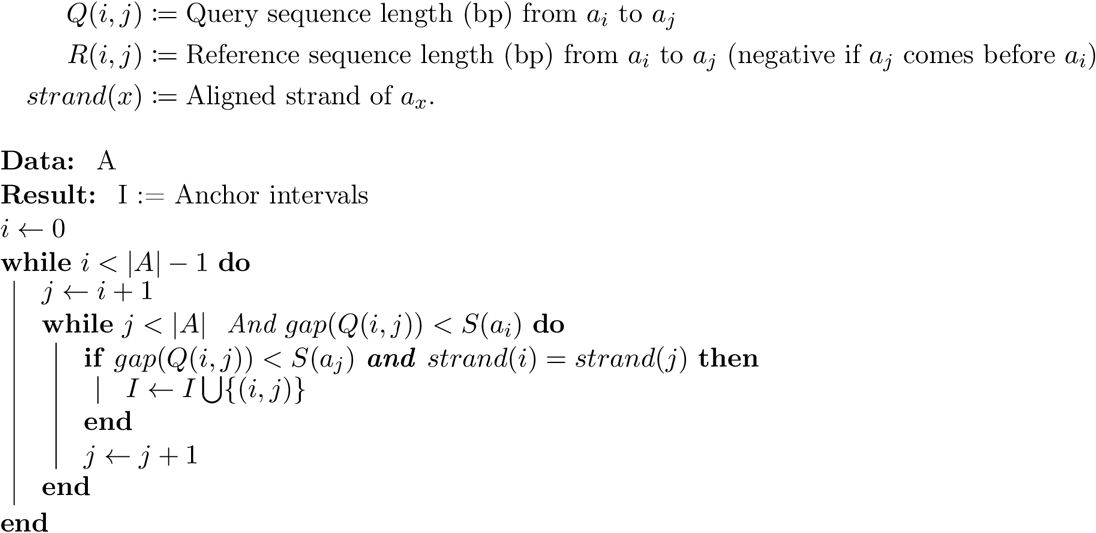

The resulting set of anchor pairs, *I*, defines intervals over the ordered set of alignments, *A*. Any element *a*_*i*_ of *A* may be an anchor or may fall between anchors in zero or more candidate intervals.

#### Eliminating alignment artifacts

PAV checks for alignment errors that can lead to false variant calls, for example, alignments through *TAF11L* repeats suggest a INVDUP-INV-DEL CSV, although the variant is more likely a balanced inversion (**Fig. 6**). For alignments that leave a reference gap (R(i,j) > 0), R(i,j) and Q(i,j) is within 85%, fewer than 10 segments are aligned, and the Q(i, j) < 1e6, PAV uses the convolution method to determine if the query sequence between anchors fits into the reference gap in forward orientation (no variant) or reverse orientation (balanced inversion). No further attempts at resolving the site are made.

**Figure 6.**
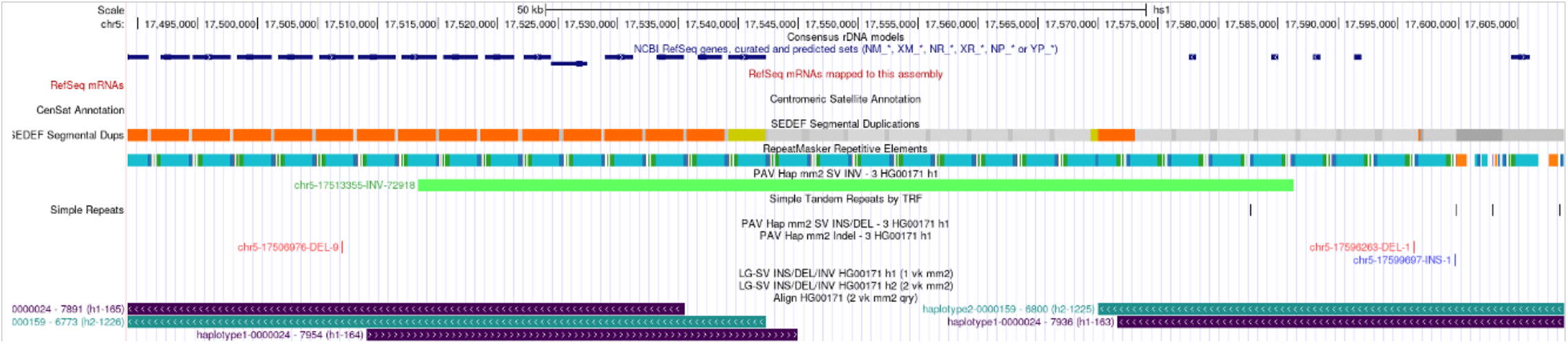
Alignment ambiguity in TAF11L repeats. Two haplotypes (violet and aqua) are aligned through TAF11L repeats on Chromosome 6 (6p15.1). One haplotype (aqua) shows alignment patterns consistent with an inverted duplication CSV (INVDUP-INV-DEL). The other haplotype (violet) shows a substitution where unaligned segment (not shown) replaces a gap in the alignment. Both represent the same variant where one aligned and one did not. PAV tests sequences between anchors to see if they fit the reference in forward orientation (no variant, likely an alignment artifact) or in reverse orientation (inversion). In this case, both alleles more likely represent an inversion and not a CSV, and PAV calls both as an inversion.

#### Variant calling between two anchors

For every anchor interval *(i, j)* in *I* that passes the alignment artifact and inversion check, PAV checks it for compatibility with different variant types, scores each one, and chooses the representation with the highest score. In the rare case of equally scored variants for a segment, simpler variants are given priority in the following order: insertions, deletions, tandem duplications, inversions, and CSVs. The requirements for each variant type and the method for assigning a score (*Vi,j*) is given below for each variant type.

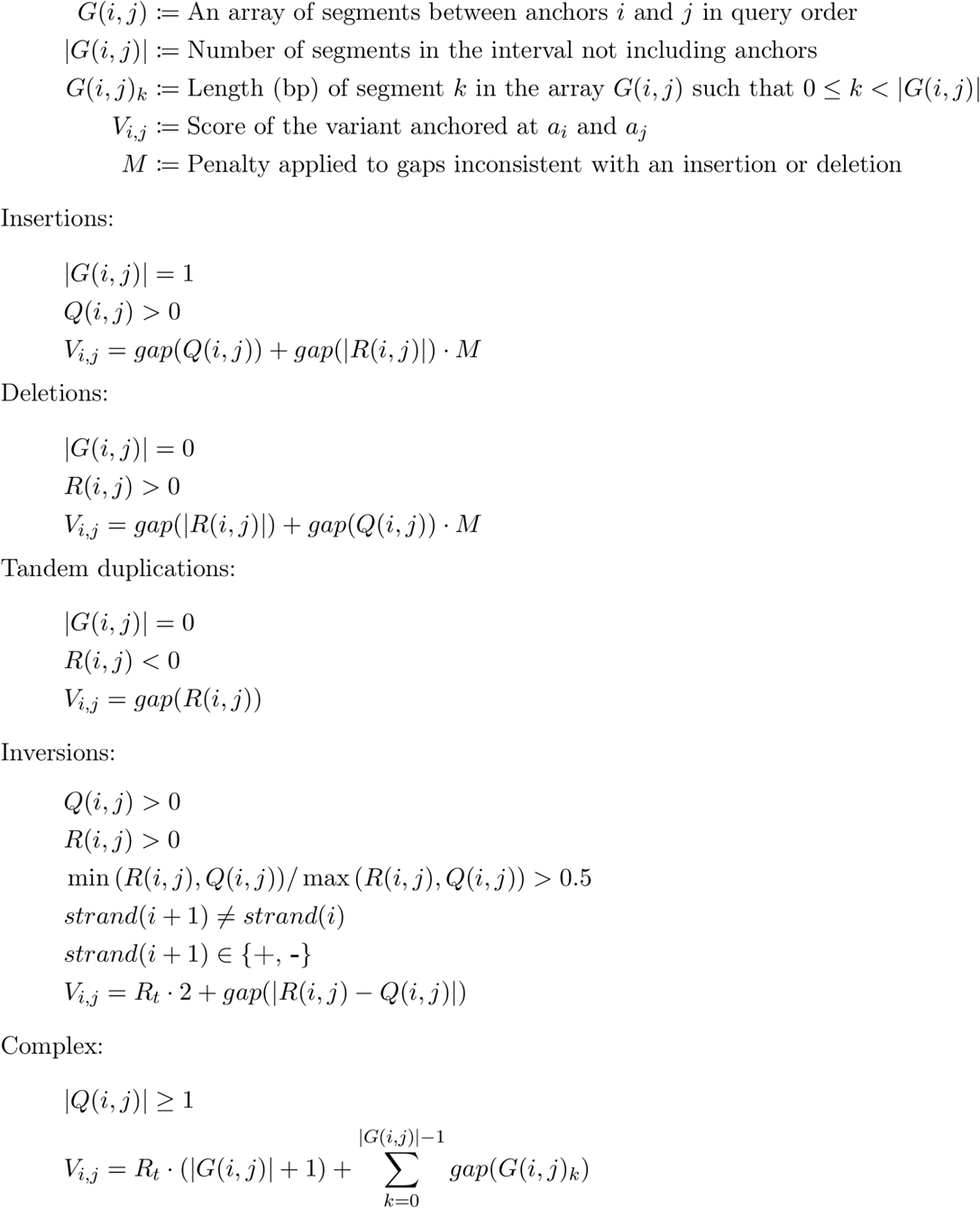

#### Choose variants across all anchor candidate pairs

A graph is constructed for each query sequence where all alignment records are vertices (V) and directed edges are created between vertices (E) such that traversing the graph strictly increases in query coordinates. Edges are composed of anchor intervals. Edges are added for all aligned nodes adjacent in the query sequence if they were not anchors. The resulting graph is a single connected directed acyclic graph (DAG) (**Fig. 7**).

**Figure 7.**
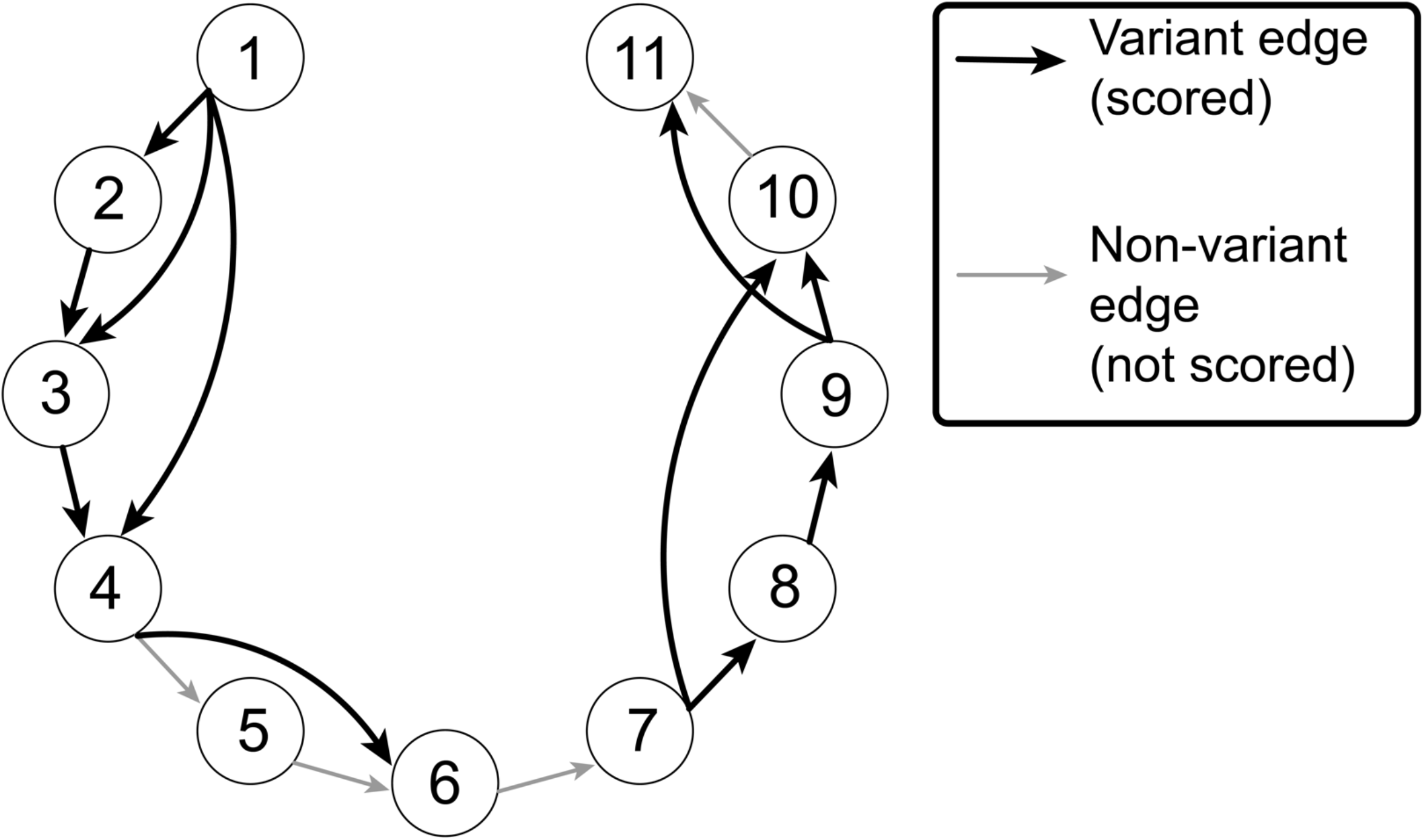
Illustration of a variant graph. Each aligned segment (numbered circle) is a vertex in the graph connected by edges and has weight equal to the alignment score computed by PAV. Each vertex (arrow) connects edges where the edges are an anchor pair (black arrows) or adjacent without a variant (gray arrows). The highest-scoring traversal of the graph determines which variants are accepted.

The weight of an edge with a variant call is the variant call score plus half the score of each anchoring alignment. Edges with no variant call has two origins, an anchor interval where no variant call could be found or an edge that was added between query-adjacent nodes. In both cases, the edge weight is computed by summing the gap penalty for each segment (excluding anchors) and adding a template switch penalty between each segment (including anchors). If the edge was an interval where no variant was found, then half of the anchor scores are added to the edge weight.

Because the resulting graph is a DAG, it is solved efficiently with the Bellman-Ford algorithm (Bellman 1958) in O(|V| + |E|) time. All variants on the optimal path through the graph are reported by PAV.

### Callset filtering

PAV CSV calls intersecting filtered loci by 50% or more were removed from the callset. Filtered regions in T2T-CHM13 were derived from the “CenSat” track in the UCSC genome browser downloaded on 2024-03-13 (https://hgdownload.soe.ucsc.edu/gbdb/hs1/censat/censat.bb). We excluded annotation types “mon” (Monomeric αSat) and “ct” (centromere transition) from this file and applied filtering using the remaining regions. For GRCh38, we utilized the “low confidence” filter we previously developed (Ebert et al. 2021).

We further filtered any variants composed of 20% or more of N’s (n-gaps) in the assembly sequence. We obtained Flagger (Liao et al. 2023) results from the HGSVC annotations (Logsdon et al. 2024) and filtered any variants composed of 10% or more of assembly sequence annotated by Flagger as non-Hap.

### Merging and intersecting CSVs

We utilize SV-Pop for merging CSVs adjusting location as the centroid of templates in the complex event for merging (variant calls after merging retain their original position). Merging parameters “nr::ro(0.5):match(0.8)” were then applied through SV-Pop to create a nonredundant callset of complex loci.

### Spectral clustering NBPF8 CSVs

The inverted sequence was extracted in reference orientation for all haplotypes with an *NBPF* CSVs, which includes both DEL-INV-DEL *NBPF8* CSVs and *NBPF9* substitution CSVs at the *NBPF8* locus. For all pairs of haplotypes, the sequence identity was determined by converting sequences to 31-mers and comparing with 31-mer depth (i.e. a 31-mer with n copies is counted n times), counting the number of intersecting 31-mers, then dividing by the number of 31-mers in the longer sequence. The table of identities was passed through spectral clustering with three clusters utilizing scikit-learn (sklearn.cluster.SpectralClustering) (Pedregosa et al. 2011).

## Notes

https://github.com/BeckLaboratory/pav

### Competing Interest Statement

The authors have declared no competing interest.

